# RBA for eukaryotic cells: foundations and theoretical developments

**DOI:** 10.1101/750182

**Authors:** Anne Goelzer, Vincent Fromion

## Abstract

Resource allocation models were recently identified as new ways to investigate cell design principles. In particular, the Resource Balance Analysis (RBA) framework is the first constraint-based modelling method capable of accurate quantitative predictions of the genome-wide resource allocation. Initially developed and validated on bacteria, the objective of this paper is to provide the mathematical fundations of the extension of the RBA framework to eukaryotic cells. We especially investigate the way to handle the cellular compartments in order to formalize eventually the functioning of organelles. It turns out that the final RBA problem for eukaryotic cells is close to the one of prokaryotic cells from a theoretical point of view. The mathematical properties that were already identified on the prokaryotic RBA framework can be easily transposed to eukaryotic cells. In particular, the eukaryotic RBA problem can be solved easily at the cell scale by Linear Programming. This paves the way to future developments of RBA models for eukaryotic cells.

## 1 Introduction

Recently, bacterial whole-cell models based on concepts of parsimonious resource allocation between cellular processes were shown to predict accurately the growth rate and a high number of cellular variables (i.e. metabolic fluxes, abundances of molecular machines including enzymes) across exponential growth conditions [3, 7, 2]. Moreover, they were also shown to reproduce complex genetic regulations without the explicit addition of gene regulations, thus reproducing observed cellular configurations with simple cost-benefit arguments with respect to the use of cellular resources [14]. Such types of models are so-called constraint-based models, and actually formalize the mathematical relationships defining the interactions and allocation of resources between the cellular processes as a set of equality and inequality convex constraints [5, 6]. Satisfying these constraints then led to a linear convex optimization problem. The underlying problem is thus tractable and solvable rapidly even at genome scale in a few seconds [1, 10].

The first genome-scale constraint-based modeling method, named Resource Balance Analysis (RBA) was developed in 2009-2011 [5, 6], where the first versions of the RBA model for the bacterium *Bacillus subtilis* were already developed and simulated. The first (and still only) biological validation of a genome-scale model integrating resource allocation was achieved in 2015 for the RBA model of *B. subtilis* [7]. In parallel, since 2013, other constraint-based modelling methods integrating some of the RBA constraints have been developed [11, 9], confirming that resource allocation is a relevant cell design principle and also an intensive research area within the systems biology community (see [4] and references therein). Currently, the resource allocation framework was used to generate resource allocation models for prokaryotes. There exists published resource allocation based models for the bacterium *B. subtilis* [5, 6, 7, 2], *Escherichia* coli [11, 9, 2] and the cyanobacteria *Synechococcus elongatus PCC 7942* [12]. Extending the resource allocation framework to eukaryotic cells is now the next step to achieve.

In this document we study the extension of the RBA framework that was originally developed in [5, 6] for bacteria, to eukaryotic cells. Here our purpose is not to develop and implement a RBA model for eukaryotic cells, but to provide the mathematical foundations of the framework, and especially:

- to handle the compartments on RBA constraints in a systematic way;
- to identify clearly the assumptions that are necessary to obtain a convex formulation of the constraints;
- to derive the constraints, and *in fine* the RBA optimization problem for eukaryotic cells.

As we will show in the sequel, under mild assumptions, the existence of compartments does not change the theoretical nature of the RBA problem, implying that the mathematical properties presented in [5, 6] are thus valid in case of eukaryotic cells. In particular, this means that the RBA optimization problem for eukaryotic cells can be efficiently solved at the genome scale. The paper is organized as follows. We first recall in Section 2 the RBA framework for prokaryotes, especially to introduce the notations. Section 2.1 presents the original problem of [7]. In Section 2.2, we extend the constraints to include the turnover rate of proteins and metabolites. Then, we investigate in Section 3.1, 3.2, 3.3 and 3.4 the consequences of compartments on RBA constraints. Finally, in Section 3.5, we present the RBA framework for eukaryotic cells.

### Notation

*A*^*T*^ refers to the transpose of the matrix *A*. 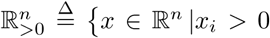 for all 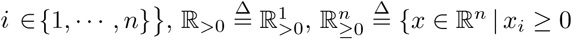 for all *i* ∈ {1, …, *n*}} and 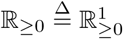

## 2 The RBA problem for prokaryotic cells: some recall

In this section, we first recall the RBA optimization problem for prokaryotes, introduce notations and also the constraints. Most of the results are extracted from [5, 6].

### 2.1 The RBA standard formulation: some recalls

A cell is composed of different cellular entities:

i. *N*_*y*_ molecular machines, that can be subdivided in *N*_*m*_ enzymes and transporters involved in the metabolic network (i.e. enzymes, transporters) 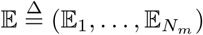 at the concentrations 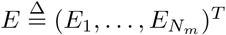 and with the fluxes 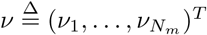; and *N_p_* macromolecular machines 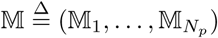 involved in non-metabolic cellular processes such as the translation apparatus, at the concentrations 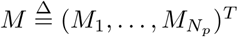;
ii. *N*_*g*_ proteins 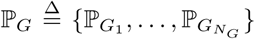 for which the cellular process to which the proteins belong is not specified. 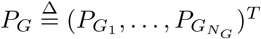 denotes the set of concentrations of ℙ_*G*_;
iii. *N*_*s*_ metabolites 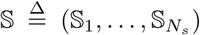 at the concentrations 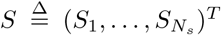. Among the set 𝕊, we distinguish a subset 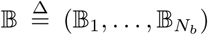 of metabolites which have fixed concentrations 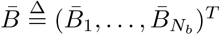.

#### Remarks

1. When a metabolic reaction can be catalyzed by two (respectively, *n >* 2) different enzymes, corresponding to isoenzymes, the metabolic reaction is duplicated (respectively, *n* times repeated), i.e., when a metabolic reaction *r* is catalyzed by 𝔼_1_ or 𝔼_2_ then we introduce two reactions *r*_1_ and *r*_2_ that are catalyzed by 𝔼_1_ and 𝔼_2_ respectively.
2. When an enzymatic complex 𝔼_*i*_ can catalyze two (respectively, *n >* 2) distinct reactions, we duplicate (respectively, repeat *n* times) the enzymatic complex: when the enzymatic complex 𝔼_*i*_ catalyzes reactions *r*_1_ and *r*_2_, then we introduce two enzymatic complexes 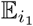 and 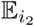 that catalyze *r*_1_ and *r*_2_ respectively. In this case, the translation process has to produce both 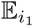 and 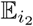.▪

The RBA optimization problem for prokaryotic cell 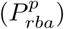 can be formalized mathematically as follows.

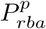: For a fixed vector of concentrations 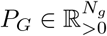, and the growth rate *µ* ≥ 0, i.e. the amount of produced biomass per cell per hour,

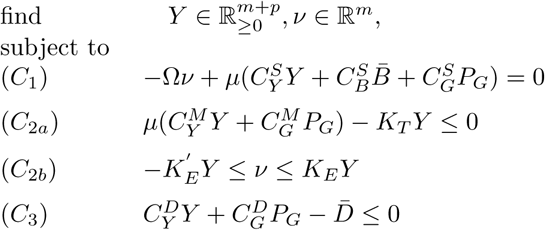

where all the inequalities are componentwise inequalities, *Y*^*T*^ ≜ (*E*^*T*^, *M*^*T*^) is the vector of concentrations of molecular machines and:

- Ω is the stoichiometry matrix of the metabolic network of size *N*_*s*_ × *N*_*m*_, where Ω_*ij*_ corresponds to the stoichiometry of metabolite 𝕊_*i*_ in the *j*-th enzymatic reaction;
- 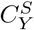 (resp. 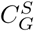) is a *N*_*s*_ × *N*_*y*_ (resp. *N*_*s*_ × *N*_*g*_) matrix where each coefficient 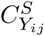 corresponds to the number of metabolite 𝕊_*i*_ consumed (or produced) for the synthesis of one machine 𝕐_*j*_ (resp.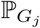); 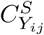 is then positive, negative or null if 𝕊_*i*_ is produced, consumed or not involved in the the synthesis of one machine 𝕐_*j*_ (resp. 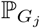);
- 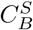 is a *N*_*s*_×*N*_*b*_ matrix where each coefficient 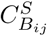 corresponds to the number of metabolite 𝕊_*i*_ consumed (or produced) for the synthesis of one 𝔹_*j*_;
- *K*_*T*_ (*K*_*E*_ and 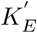, respectively) of size *N*_*p*_ × *N*_*p*_ (*N*_*m*_ × *N*_*m*_, respectively) is diagonal matrix where each coefficient 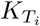 (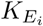 and 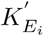, respectively) is positive and corresponds to the efficiency of the molecular machine 𝕄_*i*_, i.e. the rate of the process per amount of the catalyzing molecular machine, (the efficiency of the enzyme 𝔼_*i*_ in forward and backward sense, respectively);
- 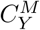 (resp. 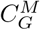) is a *N*_*p*_ × *N*_*y*_ (resp. *N*_*p*_ × *N*_*g*_) matrix where each coefficient 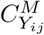 typically corresponds to the length in amino acids of the machine 𝕐_*j*_ (resp. 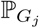). In some cases (for instance for the constraints on protein chaperoning), the length in amino acids can be multiplied by a coefficient, such as the fraction of the whole proteome that necessitates chaperoning;
- 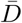 is a vector of size *N*^*c*^, where *N*^*c*^ is the number of volume and surface areas for which density contraints are considered. 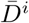 is the density of molecular entity with respect to the volume or surface area. Densities are typically expressed as a number of amino-acid residues by volume or surface area.
- 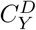 (resp.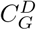) is a *N*^*c*^ × *N*_*y*_ (resp. *N*^*c*^ × *N*_*G*_) matrix where each coefficient 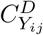 corresponds to the density of one machine 𝕐_*j*_ (resp. 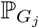) in the compartment *i*. By construction, we have one unique localization per machine.

Actually, the RBA method formalizes the mathematical relationships defining the interactions and allocation of resources between the cellular entities. All these relationships take the form of linear growth-rate dependent equalities and inequalities, and form the convex feasibility problem 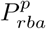[5, 6]. For cells growing in exponential phase at a growth rate *µ*,

I. the metabolic network has to produce all metabolic precursors necessary for biomass production (equalities *C*_1_ in the optimization problem);
II. the capacity of all molecular machines must be sufficient to ensure their function, i.e. to catalyze chemical conversions at a sufficient rate (inequalities *C*_2*b*_ for the enzymes and transporters, *C*_2*a*_ for the molecular machines of macromolecular processes);
III. the intracellular density of compartments and the occupancy of membranes are limited (inequalities *C*_3_);
IV. mass conservation is satisfied for all molecule types (equalities *C*_1_).

#### Remarks

1. In practice, the vector 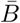 contains non-zero values only for the concentrations of macro-components such as DNA, cell wall, and lipid membranes, and for a few set of metabolites (see supplementary data of [7]).
2. In order to model reversible enzymes, we have introduced two diagonal matrices of enzyme efficiencies, i.e. *K*_*E*_ and 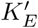, describing the constraints attached to each enzyme in both directions. Actually, when an enzyme is assumed irreversible, by convention, the constraints associated to the backward direction is taken equal to 0.
3. In [6, 7], the RBA problem that was built for *Bacillus subtilis* integrates two macromolecular processes in constraint *C*_2*a*_, the translation and chaperoning of proteins, and two density constraints, the limitation of the cytosolic density and of the membrane occupancy.
4. The RBA model can be refined by integrating for instance other molecular machines or other types of constraints such as the transcription machinery, or the protein secretion (see [7]) to cite very few.▪

To illustrate the facility to integrate new constraints, the following section presents how to integrate the degradation of molecular entities into RBA.

### 2.2 Integration of biological macromolecular turnover into RBA

The objective here is to show how to integrate the degradation of any molecular entity such as proteins, RNAs, metabolites or macromolecular machines into RBA. We restrict our developments to metabolites and proteins, other molecular entities can be handled in the same way.

#### Metabolite turnover

Some bacterial macromolecular components such as the membrane or some metabolites are damaged over time and need to be degraded and replaced. We then consider that a metabolite 𝕊_*j*_ is degraded at rate 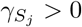. Then, the dynamics of the concentration *S*_*j*_ is given by

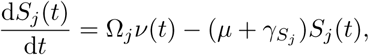

which leads in steady-state to deduce that for all the metabolites 𝔹_*j*_ in 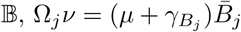.

#### Protein turnover

Actually, few of the bacterium’s proteins are systematically degraded, with the exception of a small number of proteins generally associated with regulatory or stress mechanisms. The vast majority of proteins are stable and have an average half-life of several hours (days). Eventually, the half-life of this set of stable proteins results from the fact that they are damaged over time (by temperature or specific stresses) and are typically recognized and degraded by dedicated proteases.

We then consider that a protein has a specific turnover rate 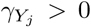. The dynamics of the concentration of protein *Y*_*j*_ is given by

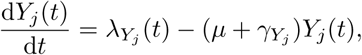

where the flux 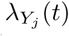 is the flux of protein (in number of molecules per unit of time and per unit of volume) produced by the ribosome. In steady-state, we thus have: 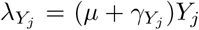. Moreover, when the protein is degraded, amino-acids are released at a rate of 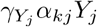, where *α*_*kj*_ is the number of the *k*-th amino acid in one protein. ATP is also consumed by the protease, at a rate of 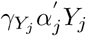 where 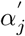 is the number of ATP consumed by the protease to degrade the protein.

##### Remark

For fast-growing bacteria such as *E. coli* or *B. subtilis*, the growth rate is usually larger than the turnover rate of most of the proteins. Therefore, protein degradation can be neglected without a significant impact on the predicted resource distribution by RBA.▪

#### Constraint on protease processes

In order to degrade proteins, the cell has to possess enough proteases. That leads to introduce a constraint on the capability of the protease to degrade proteins:

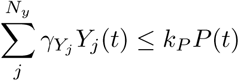

where *P*(*t*) corresponds to the concentration of proteases. This concentration is clearly a new decision variable of the RBA problem. It is integrated in the vector *Y* since the protease is a new molecular machine.

#### RBA problem for prokaryotes with protein and metabolite turnover

The set of macromolecular machines 𝕄 is now increased with the protease concentration (and *N*_*p*_ is updated in accordance). The RBA problem is now given by : for a fixed vector of concentrations 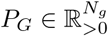, and the growth rate *µ* ≥ 0,

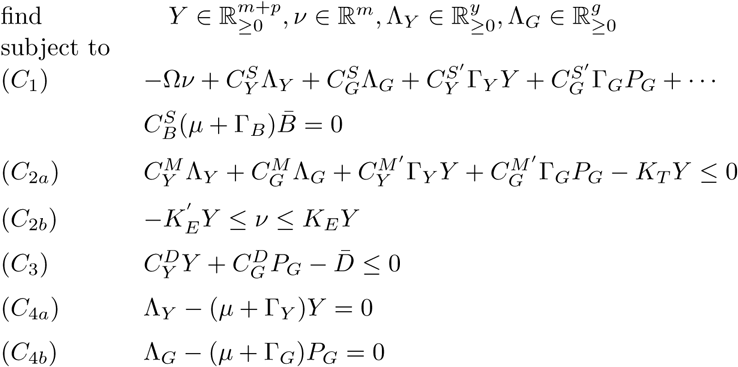

Γ_*Y*_, Γ_*G*_ are diagonal matrix of size *N*_*y*_ × *N*_*y*_, *N*_*g*_ × *N*_*g*_ respectively, and where the coefficient 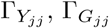 correspond to the turnover rate of 𝕐_*j*_ and 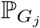 respectively. Γ_*B*_ is a *N*_*s*_ × *N*_*b*_ matrix where 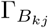 is the turnover rate of 𝔹_*j*_, and index *k* refers to the position of 𝔹_*j*_ in the set 𝕊. Moreover, 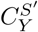 (resp.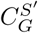) is a *N*_*s*_ × *N*_*y*_ (resp. *N*_*s*_ × *N*_*g*_) matrix where each coefficient 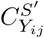 (resp. 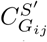) corresponds to the number of metabolite 𝕊_*i*_ consumed (or produced) during the degradation of one machine 𝕐_*j*_ (resp 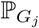).

In addition to translation and chaperoning, *C*_2*a*_ now contains the constraint on the capability of proteases to achieve protein degradation (terms 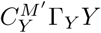 and 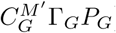), and thus the matrix 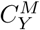 and 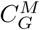 have been updated appropriately.

Moreover, constraint *C*_4*a*_ and *C*_4*b*_ correspond to the steady-state regimen of the differential equation of *Y*_*j*_(*t*) for all *j* ∈ (1 …*N*_*y*_) and on the concentration of the ℙ_*G*_ proteins.

By substituting constraints (*C*_4*a*_) and (*C*_4*b*_) in the other ones, and by noting 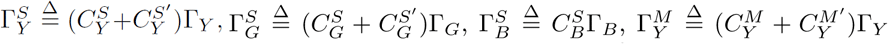, and 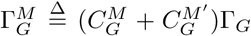, the previous optimization problem can further be aggregated as

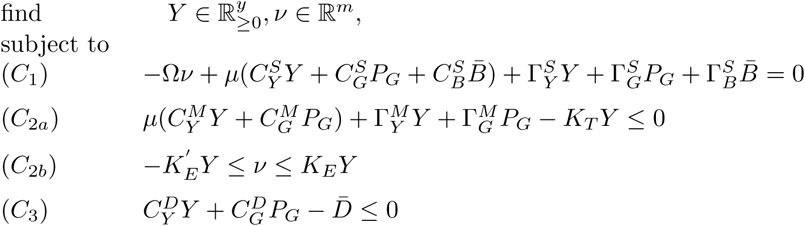

##### Remark

As in the case of protein degradation, machines that degrade cell macro-components or metabolites can also be defined and integrated into the problem in the same way.▪

## 3 RBA for eukaryotes

Eukaryotic cells are obviously much more complex than prokaryotic cells, the precise control of a large diversity of cellular processes are only partially known, even for well-established process like transcription. From a RBA perspective, we have mainly to integrate systematically in the RBA framework the presence of organelles and so to handle the fact that the metabolism, the macromolecular processes, the molecular machines and the associated constraints are now localized in different cell compartments. Actually, we notice that in genome-scale metabolic models, organelles are usually integrated only through the localization of metabolites. In the latest release of the yeast consensus model [8], the localization of some reactions is added in the reaction name as an annotation. However, the biomass reaction typically integrates the composition of the whole-cell, without distinguishing the composition of organelles. The allocation of resources to the organelles is thus fixed and constant for any condition. Hereafter, the formulation will bypass this assumption and the cell investment in the organelle is explicitly included in the resource allocation problem.

### 3.1 Modeling organelles: first steps

We define a cell as a complex system where its cytoplasm contains organelles which are described through the introduction of *N*^*com*^ compartments, with *N*^*com*^ ≥ 2 since we always assume that the cell contains at least mitochondria (see Figure 3.1). It is furthermore necessary to add the *N*^*int*^ interfaces between the different compartments. At the time *t*, the *i*-th compartment has a volume *V*^*i*^(*t*). It furthermore has interfaces with the other compartments, where each of its interfaces has a given surface area denoted 𝒜^*↔*i*^. *V*^*c*^(*t*) is the volume of the cytoplasm (or the cell^1^) at time *t* and is defined by

**Figure 1:**
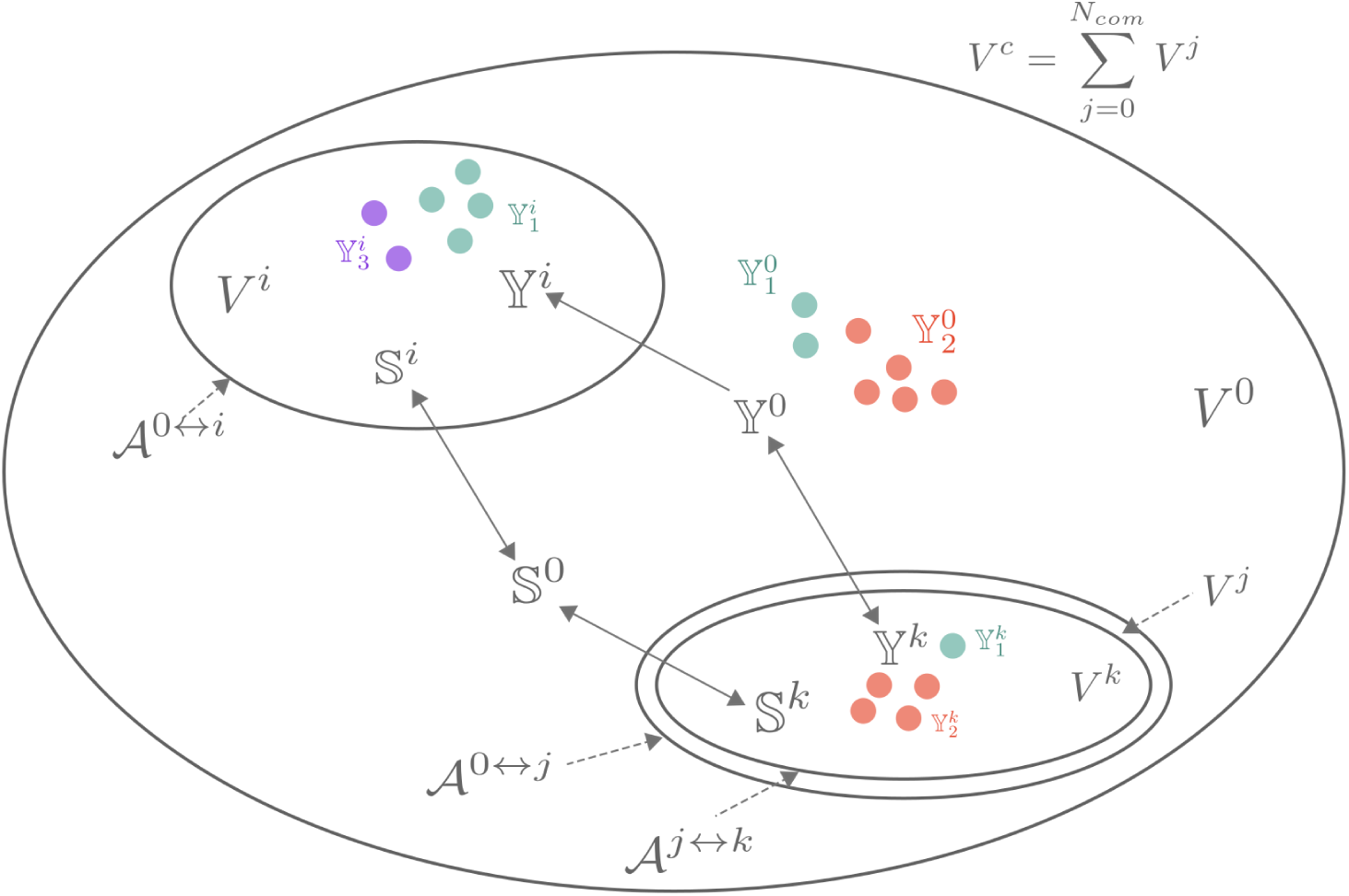
The eukaryotic cell with compartments. The cell is divided in *N*^*com*^ compartments of volume *V*^*z*^ for *z* ∈ {1,…, *N*_*com*_} and in *N*^*int*^ interfaces of surface area 𝒜^*z*^ for *z* ∈ {0 ↔ *i*,…, *j* ↔ *k*. The cell is also composed of a set of metabolites 𝕊 and of molecular machines 𝕐, that can be located in different compartments (e.g. the sets 𝕊^*k*^ and 𝕐^*k*^ for the *k*-th compartment). On this figure, we considered two organelles. The first one of volume *V*^*i*^ is composed of a single membrane of surface area 𝒜^0↔*i*^. The second one is composed of two membranes (for instance an inner and an outer membrane) of surface area 𝒜^0↔*j*^ and 𝒜^*j*↔*k*^ defining an intermembranous space of volume *V*^*j*^ and a matrix of volume *V* ^*k*^. The second organelle is thus composed of two compartments of volumes *V*^*j*^ and *V*^*k*^ associated to two interfaces of 𝒜^0↔*j*^ and 𝒜^*j*↔*k*^ respectively.

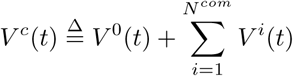

where *V*^0^(*t*) corresponds to the volume at time *t* of the part of the cell volume not contained in one of *N*^*com*^ compartments, and classically called the cytosol.

#### Remarks

1. In the sequel, the volume of the cytosol is the one which allows to define the concentration of an entity present in this part of the cell.
2. To describe a mitochondrion, two compartments must be defined: the intermembrane space and the matrix. A single interface is associated to the matrix and corresponds to the inner mitochondrial membrane. The intermembrane space has two interfaces, one with the matrix, already mentioned, and a second one with the cytosol corresponding to the outer mitochondrial membrane.▪

In order to model the cell, we then introduce a catalog of entities (defined in detail later) where it is assumed that each entity belongs to a specific compartment or/and to an interface between two compartments. The abundance of each entity is assumed to be known in each compartment and interface: 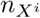 corresponds to the number of 𝕏 in the *i*-th compartment and [*X*^*i*^]^*i*^ corresponds to its concentration in the *i*-th compartment, i.e., 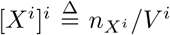. Finally, [*X*^*i*^]^*c*^ defines the concentration of 𝕏 in the *i*-th compartment with respect to the cell volume, i.e., 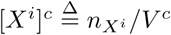. The total concentration of 𝕏 with respect to the cell volume is then given by

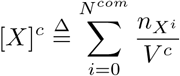

where 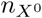 corresponds to the number of 𝕏 in the cytosol.

For an interface between the *i*-th and *j*-th compartment, the number of 𝕏 in this interface is denoted by 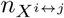.

#### Modeling the creation of an entity 𝕏 in the *i*-th compartement

We now consider that the creation of 𝕏 in the *i*-th compartment is described by this differential equation:

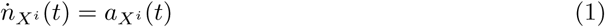

where 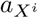 is then a quantity proportional to a rate, i.e., a number of molecules per unit of time. The evolution of the concentration of this entity in its compartment is obtained after straightforward computations and given by

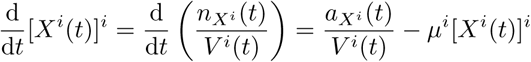

where 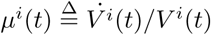.

Following the same lines, it is possible to express the evolution of 𝕏 in the *i*-th compartment through the evolution of its concentration with respect to the cell volume:

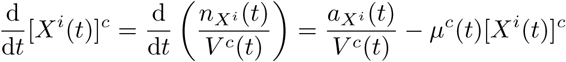

where 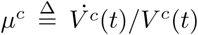. By introducing 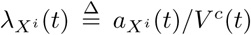, corresponding to the rate of production of 𝕏 in the *i*-th compartment per unit of cell volume, the previous differential equation is rewritten as

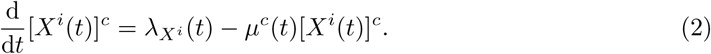

#### The density constraints attached to compartments and their interfaces

The density constraints (i.e. constraints (*C*_3_) in the RBA problem of the previous section) are related to the fact that biological entities can only occupy a part of the volume of each compartment or of the surface defining the interfaces between the different compartments.

By virtue of a formulation introduced in [5, 6], we then assume that the macromolecules present in a compartment occupy a maximal fraction of the compartment volume (respectively, surface). Thus, we introduce *β*^*i*^, a global parameter, attached to the *i*-th compartment. In the RBA framework, the coefficient *β*^*i*^ is expressed as an equivalent volume occupied by one amino acid in this compartment. This is used to express the volume or the surface occupancy of each macromolecule in terms of an equivalent number of amino acids (see [6, 7] for details). Finally, we can link the volume of the compartment to its content in macro-entities, as e.g. enzymes, chaperones, ribosomes, by this relationship:

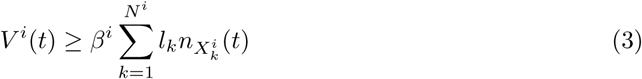

for each *i* ∈ {0, 1, …, *N*^*com*^} and where *N*^*i*^ is the number of possible (macro) entities present in the *i*-th compartment and *l*_*k*_ for each *k* ∈ {1, …, *N*^*i*^} is the total number of equivalent amino acid residues of the entity 𝕏_*k*_.

Following the same lines, the density acting on the surface area of the interface between the *i*-th and *j*-th compartment is given by *β*^*ij*^, expressed as the equivalent surface occupied by one amino acid in this interface, and this relationship:

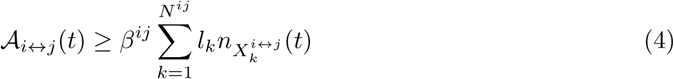

where *N*^*ij*^ is the number of possible (macro) entities present in the interface between the *i*-th and the *j*-th compartment and *l*_*k*_ for each *k* ∈{1, …, *N*^*ij*^} is the total number of amino acid residues of the entity 𝕏_*k*_.

##### Remark

More generally, the volume *V* ^*i*^ and the surface area *A*_*i*↔*j*_ of a compartment are usually linked by the so-called volume/surface ratio *σ*_*i*_. By the definition, the volume/surface ratio *σ*_*i*_ of the *i*-th compartment is given by

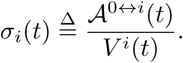

We will see in Section 3.5 how the relation between the volume and the surface area of a compartment is taken into account into RBA constraints.▪

#### A simplifying assumption

In order to simplify the presentation in the reminder part of this article, we assume that the volume of each compartment, including cytosol, saturates the density constraint over time, i.e., Inequality (3) is an equality for all compartments. This implies that the cell volume is relied to the (macro) entity content of each compartment by this simple relation:

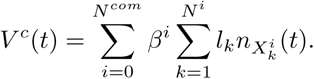

The evolution of the cell volume with respect to the time is then given by

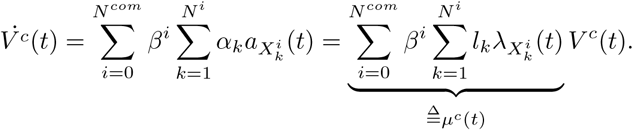

The expression of the cell volume leads to define a first constraint on the concentrations of the set of all (macro) entities present in the cell:

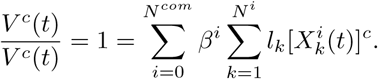

If we furthermore introduce the volume of each compartment as a fraction of the cell volume:

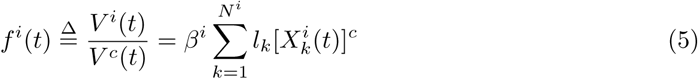

for all *i* ∈ {0, 1, …, *N*^*com*^}. We thus conclude that

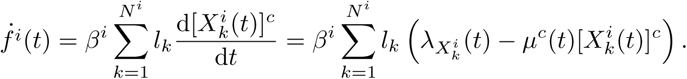

Moreover, the volume fraction *f*^*i*^ can be related to the surface area of the compartment normamized by the cell volume through Equation (4) as follows:

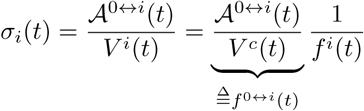

This leads to a constraint between the surface area and the volume of the compartment:

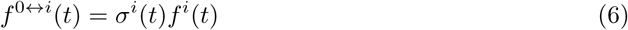

##### Remark

Other assumptions can be easily considered, in particular if the active constraints are the interface surface rather than the volume. Finally, if the volume or interface surfaces of compartments are defined differently by other mathematical expressions, it is important to verify that the associated constraints can be formulated as convex constraints.▪

### 3.2 How to handle the flux exchanges between different compartments

We now investigate how the exchange fluxes of metabolites between two compartments are computed. In order to simplify the presentation, we consider the *i*-th compartment of volume *V* ^*i*^, which has an interface with the cytosol of volume *V* ^0^. We will also first assume that the volumes of compartments (*V* ^*i*^ and *V* ^0^) are constant.

The metabolic network in the *i*-th compartment is classically described by a stoichiometric matrix Ω^*i*^ and the evolution of metabolite concentrations is described by

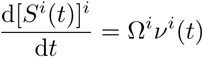

where [*S*^*i*^(*t*)]^*i*^ is the vector of metabolite concentrations (in the *i*-th compartment), *ν*^*i*^(*t*) is the vector of fluxes (expressed as a number of molecules per unit of time and per unit of volume), concatenating internal fluxes, denoted 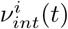, and fluxes of exchange between the compartment and the cytosol, denoted 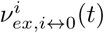. By convention, an exchange flux is positive when the molecule is imported in the compartment and negative when it is exported from it. In such a model, since all the quantities are defined per unit of volume and are attached to a specific compartment, the exchanges with another compartment have to be converted, typically as a number of molecules per unit of time, i.e.,

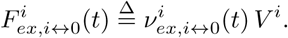

This conversion allows us to write the equation linking the exchanges between the two considered compartments as

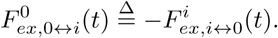

However, the previous expression has a general drawback since the fluxes are now expressed as an absolute number of molecules, and this number typically increase exponentially during a growth phase. But that is not really a problem, we have just to normalize all these numbers by *a common quantity*, chosen to be for example the volume of cell. In such a way, we then handle only ‘normalized fluxes’ through all the compartments and these fluxes can now be added or subtracted (at the level of exchange). We then introduce:

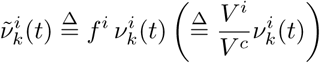

for all *k* ∈{1, …, *N*^*i*^} and all *i* ∈{0, 1, …, *N*^*com*^}. In a steady-state regime, the fluxes through metabolic networks satisfy these equality constraints:

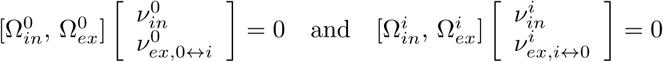

which can be rewritten with respect to the normalized fluxes as

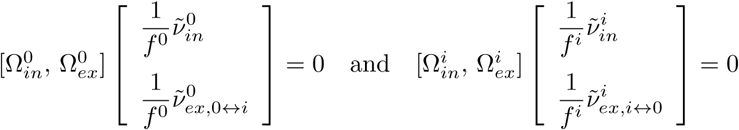

where *f* ^0^ and *f*^*i*^ are defined by Relation (5). We conclude that the interconnection of both metabolic networks in steady-state can be written as

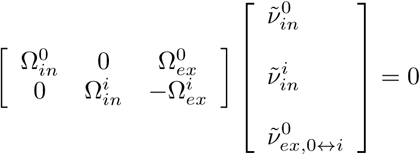

#### Compartments with time-variant volumes

The previous results are obtained when the volume of compartments is constant. We now consider a more general case. To do this, the effect of dilution due to the increasing compartment volume has to be included. This is classic and leads to the following differential equation:

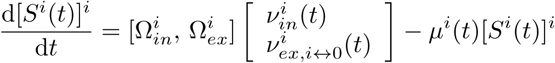

with 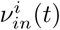 and 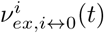 are the internal and the exchange fluxes between the *i*-th compartment and the cytosol respectively. By definition, we have

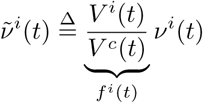

and then

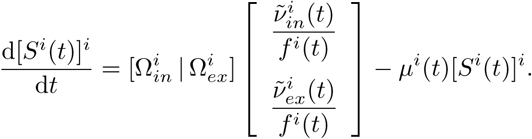

Since

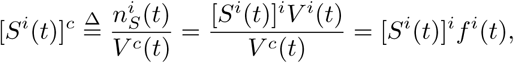

then we have

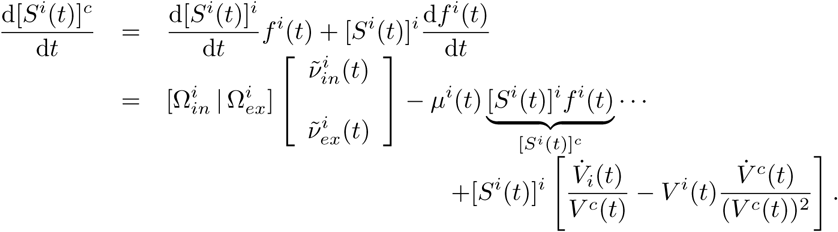

Finally, since 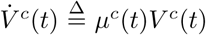 and 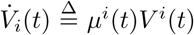, the terms depending on *µ*^*i*^(*t*) vanish, so we can rewrite the previous differential equations as

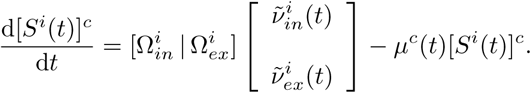

### 3.3 How to link enzyme activities in the different compartments with each other

Typically, the activity of the *k*-th enzyme localized in the *i*-th compartment is described by

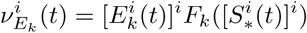

where 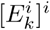 is the concentration of the *k*-th enzyme in the *i*-th compartment, 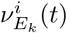 is the flux through the enzyme expressed as a number of molecules per unit of time and per unit of volume. *F*_*k*_ is the nonlinear characteristic of the enzyme activity which is a function of a set of metabolite concentrations in the *i*-th compartment, i.e., 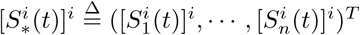. The notation 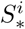 will refer in the sequel to a set of metabolite concentrations in the *i*-th compartment.

In the RBA framework, the maximal flux through the *k*-th enzyme is constrained by its concentration through this simple linear inequality:

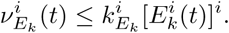

This last inequality can be rewritten in an equivalent way as an inequality between a normalized flux and the concentration of the enzyme with respect to the cell volume:

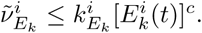

### 3.4 How to handle protein production in different compartments

#### Protein production

We assume that a protein 𝕐 is localized in the *i*-th compartment, has a given concentration [*Y* ^*i*^]^*i*^, and that 𝕐 is produced by the machine^2^ R^0^ that is located in the cytosol. We also assume that the volumes of compartments and of the cytosol are constant. So *V* ^*i*^, *f*^*i*^, *V* ^0^, *f* ^0^ and *V* ^*c*^ are constant.

The total flow of proteins (in number of molecules per unit of time) produced by the 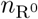 machines is 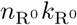, where 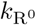 is the efficiency of R^0^ (in a number of amino acid per unit of time). Actually, like for an enzyme, the efficiency 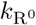 can be a nonlinear function that depends on the concentration of a set of metabolites (located in the same compartment than R0), 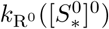. The machines produce a set of proteins that can be located in the same compartment or translocated in another one.

Let us recall that the creation (in number per unit of time) of 𝕐in the *i*-th compartment is given by 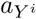 (see Relation (1)). So the capability of machine R^0^ has to be sufficient to produce 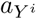, and, by extension, all the proteins that R^0^ must produce. So we have:

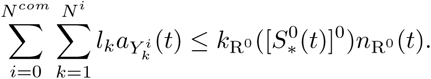

In the previous equation, *l*_*k*_ is the length in amino acids of the protein 𝕐_*k*_. In order to alleviate the notations, we assume that R^0^ produces all proteins of the cell. The formula can be extended directly to a subset of proteins by defining a subset of proteins per compartment. We can finally normalize the previous inequality by *V* ^*c*^ on both sides in order to deduce this inequality:

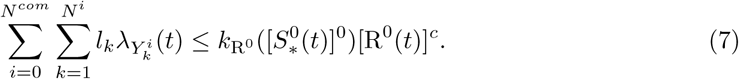

##### Remark

Normally, the constraints in the compartment 0 is expressed with respect to the concentration in the compartment 0:

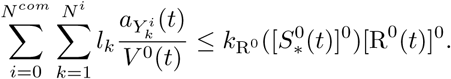

However, since 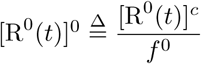, and 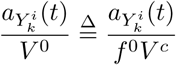, the two inequalities are equivalent.▪

#### Impact of protein production on metabolites

The protein production consumes metabolites such as charged-tRNAs, GTP and releases others such as uncharged-tRNAs, GDP and phosphate. In the case of translocation processes, these metabolites can further be located in different compartments.

We first consider the case where all metabolites that are produced/consumed are located in the cytosol. To this end, we consider a metabolite 𝕊_*k*_ located in the cytosol, and that 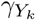 copies of 𝕊_*k*_ are consumed for the synthesis of the protein 𝕐. The flux 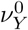 (in number of molecules per unit of time and per unit of volume) of 𝕊 *k* consumed by R^0^ to produce 𝕐 in the cytosol is equal to 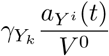. So we have:

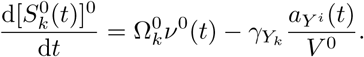

##### Remarks

1. If the metabolite is produced, then the term 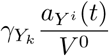 is added and not substracted in the equation.
2. If the protein 𝕐 is produced in the cytosol, and then remains in the cytosol instead of being translocated in another compartment, the flux 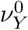 is equal to 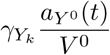.▪

By using the same notation than the previous section, we have:

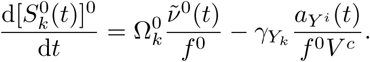

And for all proteins that are produced by R^0^ in the cytosol, we have:

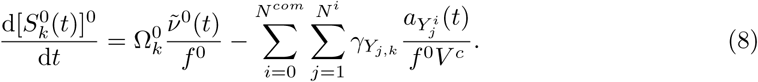

#### Steady-state regime

In steady-state, from Equation (2), Inequality (7) becomes

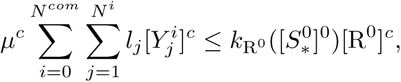

and Equality (8) becomes:

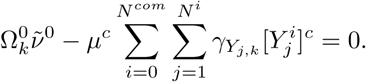

#### Compartments with time-varying volumes

Now let us consider that the volumes are time-varying. The dynamics of 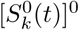 with respect to time is given by

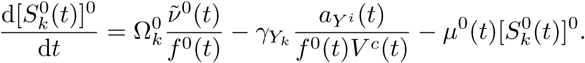

Since we have 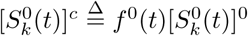 then

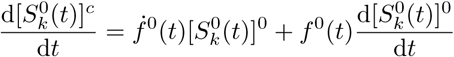

and by summing on all proteins:

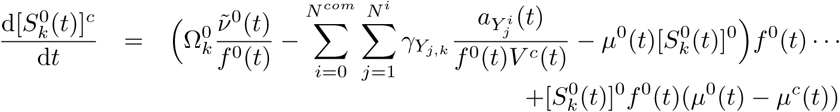

which finally leads to deduce that

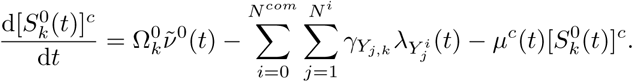

#### Protein translocation

We now consider the translocation process. A molecular machine, the translocon, translocates proteins from one compartment to another one, and usually consumes and produces metabolites that can be located in both compartments. One metabolite (such as protons) can also be translocated with the protein. In this paragraph, we establish the formulation of constraints related to the translocon capability, and also on the associated metabolites.

We consider (*i*) 3 metabolites 𝕊_1_, 𝕊_2_ and 𝕊_3_. 𝕊_1_ is located in the cytosol, 𝕊_2_ in the *i*-th compartment, and 𝕊_3_ is translocated between compartments; (*ii*) the translocon 𝕋 that is localized in the membrane; and (*iii*) a protein 𝕐 that is translocated. The membrane is considered as the 0 ↔ *i* interface. Moreover, we consider that all volumes are constant.

As in the previous sections, we have a number of translocon 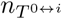 that catalyzes the translocation of *n*_*Y*_ copies of 𝕐 from the cytosol to the *i*-th compartment. In most of organisms, such translocation steps are irreversible. We thus have one translocon to translocate proteins from the cytosol to the *i*-th compartment (e.g. Tim/Tom, Tic/Toc machines). What count in our model will be the active protein 𝕐 in the *i*-th compartment. We will thus note 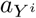 the number of 𝕐 that will be translocated per unit of time and that will be present finally in the *i*-th compartment.

The capability of the translocon (in number per unit of time) must be sufficient to translocate the flow of proteins 𝕐 (also expressed in unit of time). So we have the constraint:

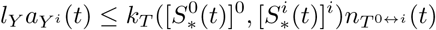

with 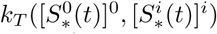 the efficiency of the translocon (in number of amino acids residues per unit of time), depending on a set of metabolites that can be located in both compartments, and *l*_*Y*_ the length of amino acid residues of 𝕐. We can normalize the previous inequality by *V* ^*c*^ and obtained

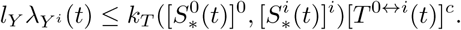

And for all proteins that are translocated by 𝕋, we have:

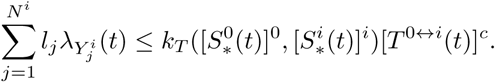

#### Impact of translocation on metabolites

For 𝕊_1_ and 𝕊_2_, we have the same type of relations than in the previous paragraph. The flux (in number of molecules per unit of volume per unit of time) of 𝕊_1_ (located in the cytosol) consumed by 𝕋 to translocate 𝕐 is equal to 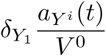, where 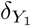 is the number of 𝕊_1_ consumed per translocated protein. So for all proteins:

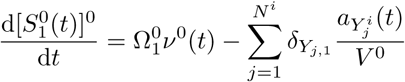

where 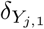 is the number of 𝕊_1_ consumed to translocate *Y*_*j*_. And for 𝕊_2_:

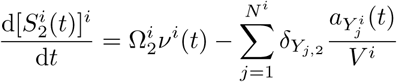

where 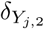 is the number of 𝕊_2_ consumed to translocate *Y*_*j*_.

By normalizing by *V* ^*c*^ instead and by using the same notations than before, we have:

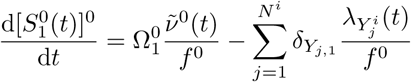

and:

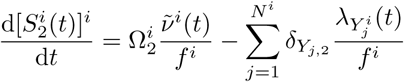

Let us now consider 𝕊_3_ and the two concentrations in the cytosol and in the *i*-th compartment. The flux of 𝕊_3_ (in number of molecules per unit of time) that crosses the interface 0 ↔ *i* (from the cytosol to the *i*-th compartment) is equal to:

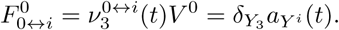

By the mass conservation, the flux of 𝕊_3_ that enters is equal to 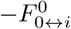. In the cytosol, the dynamics of the concentration of 𝕊_3_ when the protein 𝕐 is translocated, is given by

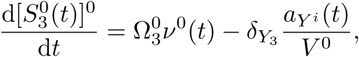

where 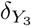 is the number of 𝕊_3_ that is translocated together with 𝕐. For the concentration of 𝕊_3_ in the *i*-th compartment, we have:

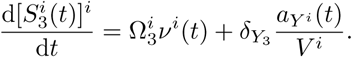

By using the same transformation than the previous section, we finally obtain:

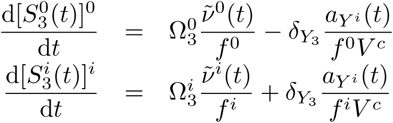

For all proteins that are translocated, we have:

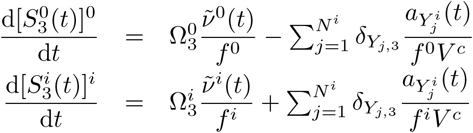

#### Steady state regimen

In steady state, we thus have

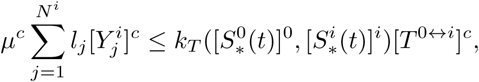

and for the metabolites 𝕊_1_, 𝕊_2_ and 𝕊_3_, these following constraints:

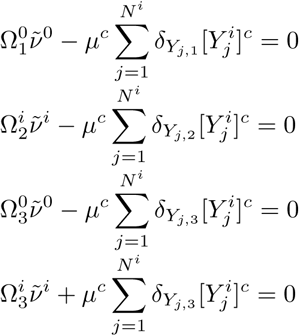

#### Compartments with time-variant volumes

For the general case, we follow the same lines than in the previous section, and obtain for 𝕊_2_ for instance:

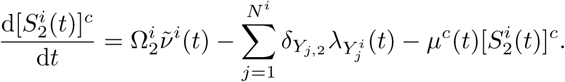

### 3.5 RBA problem for the *i*-th compartment

In this section, we write the RBA constraints for the *i*-th compartment of an eukaryotic cell. As we said previously, we consider hereafter the concentration of molecular species with respect to the volume of the cell (e.g. 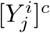 in Section 3.1). However, for the sake of readability, we will omit the brackets in the notation of concentrations in the sequel. The notation 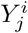 will thus refer to 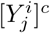, the concentration of the machine 𝕐_*j*_ localized in the *i*-th compartment with respect to the cell volume. The notation 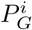 will refer to the concentration of the ℙ_*G*_ proteins in the *i*-th compartment 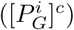. We also considered the vector of normalized flux 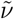. Moreover, the concentration of metabolites 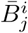 will refer to 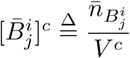, where 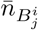 is constant and given. Again, for the sake of readibility, we will not integrate the turnover rate of molecular entities.

Thereafter, we consider the most common case: an organelle that is isolated from the cytosol by a lipidic membrane. We will consider that some metabolic and macromolecular processes occur within the organelle, and that some molecular entities (e.g. metabolites, proteins, to cite a few) are produced in the cytosol and imported into the organelle. Even if the constraints will be formulated for this type of organelle, the formulation is generic enough to describe the exchanges between any compartments.

Let us consider the *i*-th compartment and its interface with the cytosol 0 ↔ *i*. By convention, the fluxes that enter within the compartments are positive, and the ones that are excreted are negative. Let us consider the 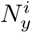 machines present in the *i*-th compartment of concentrations *Y* ^*i*^ ≥ 0, and being synthetized at rate 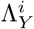 Among the 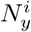 machines, we have 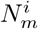 enzymes and the vector of their metabolic fluxes 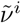. Moreover, we will further specify 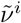 with the internal metabolic fluxes 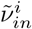 and the ones of exchange through the interface 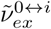.

#### Metabolic and capacity constraints

The constraints attached to the metabolic network, and to the production of molecular machines for compartment *i* are:

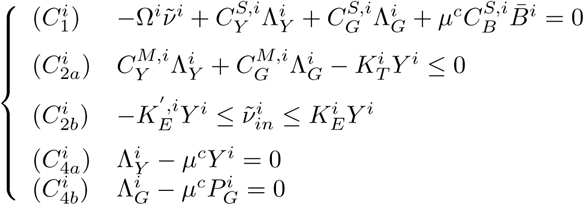

where

- 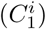 describes the mass conservation for the set of metabolites localized in the *i*-th compartment (𝕊^*i*^). Moreover, we consider that a subset of metabolites also has a fixed concentration 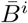 (also given with respect to the cell volume);
- 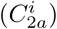 describes the capability constraints of the molecular machines localized in the *i*-th compartment;
- 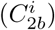 describes the capability constraints of the enzymes localized in the *i*-th compartment. Constraints on transporter are defined in the constraints related to the compartment interface.
- 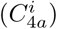 and 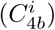 are the constraints of steady-state regimen for the dynamics of the concentration of molecular machine and ℙ_*G*_ localized in the *i*-th compartment.

In all constraints, the matrixes 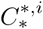 contains the same type of coefficients than the ones in Section 2.1. Only the structure of the matrices changes with respect to the type of process and to the molecular machines that are present in the *i*-th compartment and in the *j*-th constraint. For instance, if the *i*-th compartment is the mitochondria, then since translation of some mitochondrial proteins occurs only in the mitochondria, then 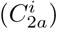 contains a constraint of the capability of the mitochondrial translation process. Therefore, we need to identify the proteins that are only translated in the mitochondria to build the constraints. If the vector 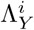 is organized along the machines that are produced in or out the mitochondria 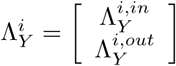 then 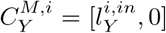 where 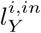 contains the length in amino acids of mitochondrial molecular machines produced in the mitochondria.

We also chose to put the constraints 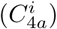 for all molecular machines of the *i*-th compartment whatever is their location of production for readability purpose.

##### Remark

As mentionned previously, the set 𝔹 of metabolites having fixed concentrations usually contains building blocks of macro-components such as DNA, cell wall or membrane, and thus in particular the building blocks for the membrane of organelles (i.e. the interface of compartment). The term 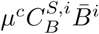 typically corresponds to the flux necessary to maintain the concentration *B*^*i*^ constant and equal to 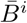 at growth rate *µ*^*c*^. If we assume that the set 𝔹^i^contains at least one metabolite 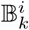 for the synthesis of any interface (for instance the interface 0 ↔ *i*), then we have:

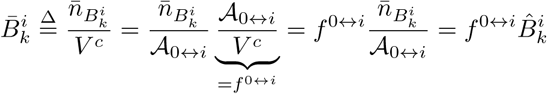

where 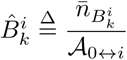 corresponds to the number of 𝔹_*k*_ per unit of surface area.

In this this case, constraint 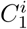 is rewritten as:

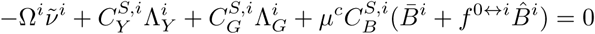

For the interface 0 ↔ *i*, we have:▪

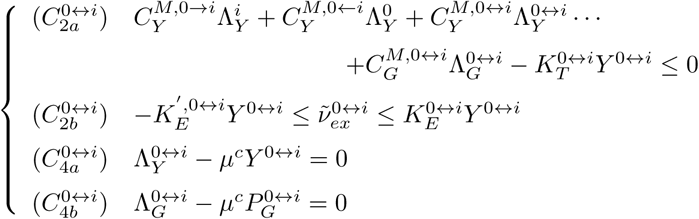

Here:

- the term 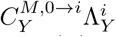 corresponds to the flux of machines that are (*i*) active in the *i*-th compartment, (*ii*) produced in the cytosol and (*iii*) translocated through the interface.
- The term 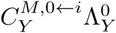 corresponds to the flux of machines that are (*i*) active in the cytosol, (*ii*) produced in *i*-th compartment and (*iii*) translocated through the interface^4^.
- The term 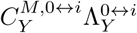 corresponds to the flux of the machines localized in the interface 0 ↔ *i* and that needs a secretion machine to be inserted in the membrane.

The vector of machines *Y* ^0↔*i*^ typically regroups the systems of import and export (if any) of proteins (e.g. TIM/TOM or TIC/TOC machine), and secretion machines to insert the membranous machines in the interface of the compartment. Only one global secretion machine could be integrated, as well as several machines (e.g. insertion of machines produced in compartment *i*, or in compartment 0) depending the level of details that we want to consider.

##### Remark

Often in eukaryotes, a molecular machine localized in the *i*-th compartment 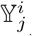 is composed of subunits that are synthetized in different compartments; that thus need to be translocated. This is well represented in the RBA framework through coefficients 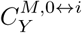. In that case, this coefficient will be equal to the length in amino acid residues of the subunits of 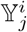 that need to be translocated.▪

#### Volume constraints

It remains to formalize the constraints on the volumes of compartments in steady-state. Some organelles such as the peroxisome, the mitochondria, the chloroplast were ancestrally bacteria. Their internal density is higher than the cytosol, and we could assume that their principle of growth follows the one of prokaryotic cells: the density of the compartment should be kept constant, and thus the increase of volume is driven by an increase in mass. Alternatively, the constraints could arise from the membrane occupancy. In this case, the increase of volume is driven by an increase in the mass of membranous proteins.

Following Section 3.1, a compartment satisfying the density constraint has this following volume:

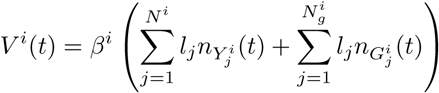

which can be normalized with respect to the cell volume leading to obtain this steady-state regimen:

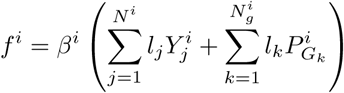

where *l*_*j*_ and *l*_*k*_ are the equivalent length in amino acid residues of the machine 𝕐_*j*_ and protein 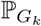 respectively, and *β*^i^ is the volume occupied by one amino acid in the *i*-th compartment. The term 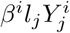 then correspond to the fraction of the volume of the cell occupied by the machine 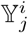.

##### Remark

*β*^*i*^ would typically correspond to 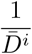 in the RBA for prokaryotic cells.▪

Moreover, we also have the constraint between the volume and the surface area of the compartment (Equality (6)):

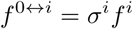

We could for instance consider a fixed surface/volume ratio σ^*i*^, so that when the volume of the compartment is higher, so is the surface.

Finally, the user should decide which constraints *f*^*i*^ or *f* ^0↔*i*^ is the active one:

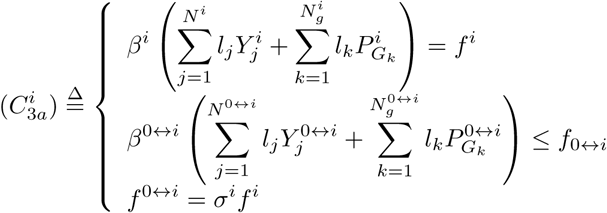

or

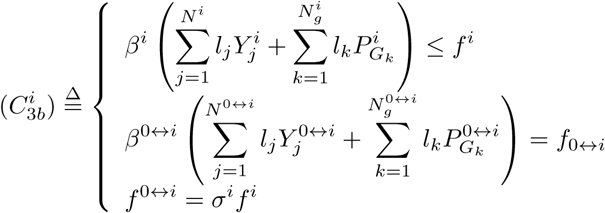

Mitochondria and the thylakoid could typically be modeled by 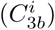 peroxisome by while chloroplast and peroxisome by 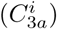.

Moreover, we could also add constraints on the minimal and maximal volume occupied by the compartment such as

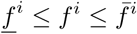

where 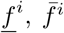 are the minimal and maximal ratio of the *i*-th compartment with respect to the cell volume respectively. Having 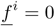 means that the compartment may be absent.

Finally, the volume of the cytosol is defined as the volume of the cell minus the volume of all organelles. In addition, the volume occupied by all machines within the cytosol must be compatible with the volume of the cytosol:

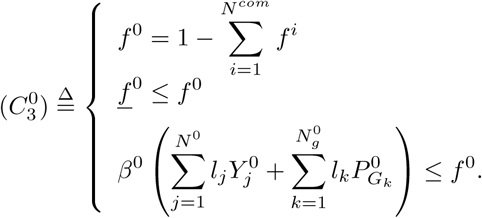

##### Remark

For complex structures of organelles, such as mitochondria or chloroplast, we expect that other constraints between volumes or surfaces should be added in order to guarantee the structure of the organelle. For instance, for mitochondria, the volume of the membranous interspace between the outer and the inner membrane and of the mitochondrial matrix should be linked such as

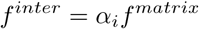

where *α*_*i*_ is the ratio between both volumes. Thus, if one of the interface or one of the volume is increasing, so are the others.

However, at this stage, the way to tackle the structure of organelles is more a modelling issue, and is thus up to the modeller. The modeler can describe the structure he wants, as soon as the mathematical formulation remains convex.▪

#### Formulation of the constraints for the *i*-th compartment

To summarize, we write now the set of constraints 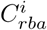 attached to the *i*-th compartment, composed of the interior and its interface by

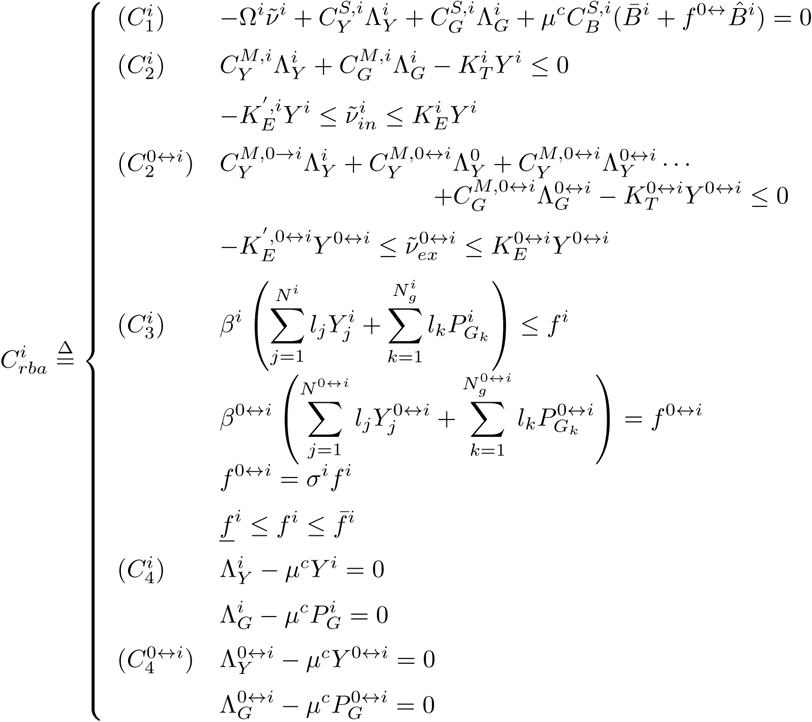

Here we assume that the constraints on volume follows the case 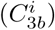. For the other type of organelles and for the cytosol, only the 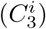 constraints need to be substituted by 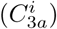 and 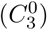 respectively.

In 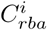, the decision variables are:

- the flux vector 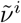 of size 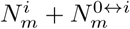
- the concentration of molecular machines *Y* ^*i*^ and *Y* ^0↔*i*^ of size 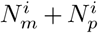 and 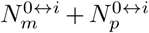 respectively
- the synthesis rates of molecular machines (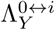 and 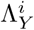 of same size as *Y*^*i*^ and *Y* ^0↔*i*^) and of ℙ_*G*_ proteins (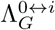 and 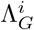 of size 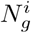 and 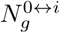 respectively)
- the volume *f*^*i*^ and *f* ^0↔*i*^.

The previous contraints can be further aggregated by introducing 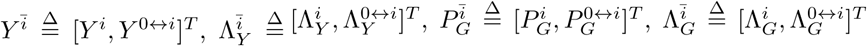, and by rewriting constraint *C*_*3*_ under a matrix form:

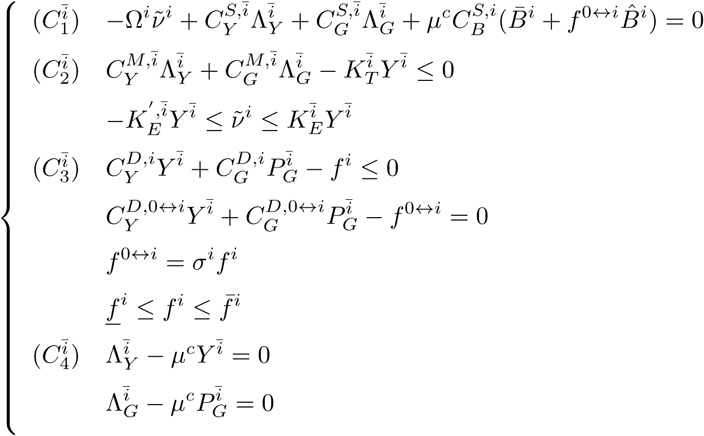

and where all matrices 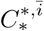 are deduced by straightforward manipulations.

Finally, if we substitute constraints 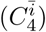 in the other ones, we obtain:

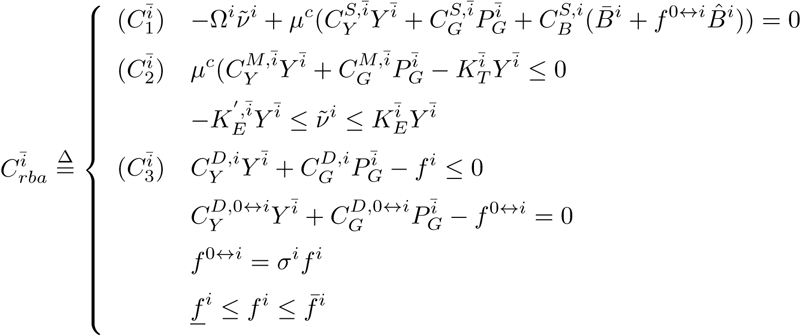

The previous formulation is thus very close to the RBA formulation of prokaryotic cells of Section 2.1. Only constraints *C*_3_ differ by having the normalized volume and surface area of the compartment as decision variables, and this, in order to adjust the volume of the organelles to the need of the cell. Morever, all constraints are linear with respect to the decision variables.

### 3.6 Formulation of the RBA problem for eukaryotic cells

The consequences of the integration of compartments on RBA have been explored in the previous paragraphs. Clearly, by introducing the normalization of all species by the cell volume, the formulation of the RBA problem for eukaryotic cells is very close to the one for prokaryotic cells. We give in this section the RBA optimization problem, 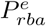, for the eukaryotic cell.

Let us introduce:

i. the set of *N*_*com*_ cellular compartments indexed by 𝕀_*V*_ = {0, …, *N*_*com*_}, and the set of *N*_*int*_ interfaces indexed by 𝕀_*A*_ = {*e* ↔ 0, 0 ↔ *i*, …, *k* ↔ *j*}5. *The set* 𝕀_*A*_ contains the indexes of all possible interfaces between compartments. We note *N*_*c*_ = *N*_*com*_ + 1 + *N*_*int*_.
ii. *Y* the vector of concentrations of all molecular machines of size *N*_*y*_ (which includes *N*_*m*_ enzymes), present in all compartments and interfaces.
iii. the flux vector *ν* of size *N*_*m*_ and associated to the *N*_*m*_ enzymes;
iv. *P*_*G*_ the vector of concentration of the ℙ_*G*_ proteins of size *N*_*g*_ present in all compartments and interfaces.
v. *f*_*V*_ is the vector of the normalized volume of compartments with respect to the total cell volume: 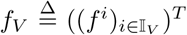. Moreover, *f*_*A*_ is the vector of normalized surface area of compartments with respect to the total cell volume: 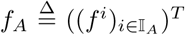. Finally, *f* is the concatenation of *f*_*V*_ and *f*_*A*_.

The RBA optimization problem can be formalized as follows.

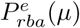: For a fixed vector of concentrations 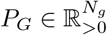, and the growth rate *µ* ≥ 0 of the cell,

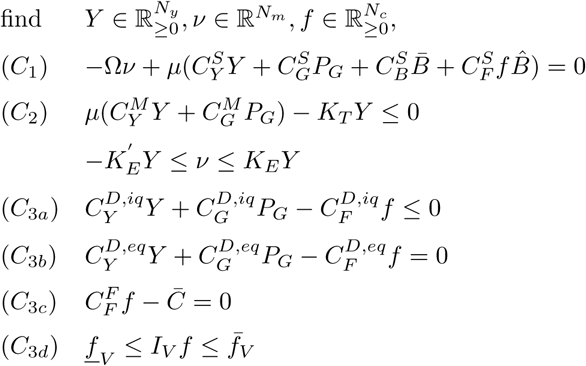

where:

- (*C*_1_) and (*C*_2_) have the same meaning than the RBA problem for prokaryotic cells
- (*C*_3*a*_) include the volume or surface constraints that may not be saturated. The contraints on the maximal density of the cytosol is typically included here.
- (*C*_3*b*_) contain the volume or surface constraints that are assumed to be always saturated
- (*C*_3*c*_) contain all the constraints between the surface and the volume of one compartment (e.g. *f*^0↔*i*^ = *σ*^*i*^ *f*^*i*^), additional constraints between volumes or surfaces for complex structures of compartments, and the equality constraint determining 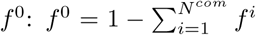. 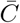 is a vector having 1 only for the constraint on *f*^0^ and zero otherwise.
- (*C*_3*d*_) regroup the constraints on the minimal and maximal volume of compartments with respect to the total cell volume. The vector *<u>f</u>*_*V*_ and 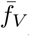 contain the minimal and maximal normalized volumes of compartments respectively, and the matrix *I*_*V*_ is such that *f*_*V*_ = *I*_*V*_ *f*.

The RBA problem for eukaryotic cells is a Linear programming optimization problem, and can thus be solved efficiently at the cell scale. Compared to the RBA problem for prokaryotic cells, the set of *C*_1_ and *C*_2_ constraints are similar, with the same matrices of coefficients and parameters 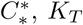 and 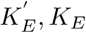. It means that this part of the optimization problem can be built exactly as in the RBA problem for prokaryotes. Only the values of efficiency coefficients 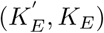 of processes could be dependent on the compartment, since they may be modulated by the internal metabolite concentrations of compartments (see Section 3.3 and 3.4).

## 4 Conclusion

In this paper, we investigated how the RBA framework can be extended to eukaryotic cells. Interestingly, it turns out that the RBA framework for eukaryotic cells remains very close to the one for prokayotic cells. More importantly, the RBA problem for eukaryotic cells for a fixed growth rate is a Linear Programming (LP) problem, and can thus be solved efficiently at the cell scale [1, 10].

Recently, we developed RBApy, a python package for the generation and simulation of RBA models for prokaryotic cells [2]. Because of the similarity between optimization problems, the extension of RBApy to eukaryotic cells should require only a few changes, mainly at the level of density constraints. Only a few additional parameters are required, and they are mainly related to the structure of organelles (such as the surface/volume ratios). Therefore, we expect no major theoretical or operational difficulty at developing RBA models for eukaryotic cells. Running RBA for an eukaryotic cell depends on the number of decision variables and of constraints. In steady-state, the associated optimization problem should be of reasonable size, enabling its efficient resolution.

In practice, developing RBA models for eukaryotic cells strongly relies on the level of knowledge on the functioning or biological processes. The framework is flexible enough to integrate a high level of details, such as all the different chaperoning systems and their targets in mitochondria, if the knowledge is available and if the modeler want to include a high level of details. Or, in contrast, the RBA model can include only an aggregated description of a biological process. Therefore, the final RBA model will result from a compromise between the level of knowledge on the functioning of the organism, the resulting size of the optimization problem, and the choice of the modeler.

RBA model parameters can be estimated from data, and even from existing published data. When we developed the RBA model of *Escherichia coli* in [2], we used existing fluxomics and quantitative proteomic datasets [15, 13] to estimate parameters, and obtained good quantitative predictions. Currently, the real challenge is the biological validation of RBA models of eukaryotic cells as we did for bacteria a few years ago in [7]: to have (or to generate) suitable datasets to learn the parameters in different growth conditions and how they evolve with respect to growth rate, in order to finally perform predictions using varying growth-rate dependent parameters. This will be achieved in a near future, as well as the update of the RBApy package to automatically generates RBA models for eukaryotic cells.

## Acknowledgments

We thank O. Bodeit and A. Bulović for their critical comments on this manuscript.

## Footnotes

1 With a slight abuse of denomination.

2 Corresponding to the translation apparatus and its associated accessory proteins.

3 Again, this is used to simplify the formulation of the RBA problem and more general modeling is possible (see [6, 7] for details).

4 Again, we introduce this term to be as generic as possible. However, the export of proteins may not occur in all organelles, so this term could vanish.

5 here *e* denotes the index of the compartment associated to the extracellular medium

## References

[1] A. Ben-Tal and A. Nemirovski. Lectures on modern convex optimization: analysis, algorithms, and engineering applications. MPS/SIAM Series on Optimization, 2001.

[2] A. Bulović, S. Fischer, M. Dinh, F. Golib, W. Liebermeister, C. Poirier, L. Tournier, E. Klipp, V. Fromion, and A. Goelzer. Automated generation of bacterial resource allocation models. Metabolic engineering, 2019.

[3] A. Goelzer and V. Fromion. Bacterial growth rate reflects a bottleneck in resource allocation. Biochemical Biophysical Acta, 1810(10):978–988, 2011.

[4] A. Goelzer and V. Fromion. Resource allocation in living organisms. Biochemical Society Transactions, 45(4):945–952, 2017.

[5] A. Goelzer, V. Fromion, and G. Scorletti. Cell design in bacteria as a convex optimization problem. Proceedings of the 48th IEEE Conference on Decision and Control, pages 4517–4522, December 2009.

[6] A. Goelzer, V. Fromion, and G. Scorletti. Cell design in bacteria as a convex optimization problem. Automatica, 47(6):1210–1218, 2011.

[7] A. Goelzer, J. Muntel, V. Chubukov, M. Jules, E. Prestel, R. Nolker, M. Mariadassou, S. Aymerich, M. Hecker, P. Noirot, D. Becher, and V. Fromion. Quantitative prediction of genome-wide resource allocation in bacteria. Metabolic Engineering, 32:232–243, 2015.

[8] H. Lu, F. Li, B. J. Sánchez, Z. Zhu, G. Li, I. Domenzain, S. Marcišauskas, P. M. Anton, D. Lappa, C. Lieven, et al. A consensus *S. cerevisiae* metabolic model yeast8 and its ecosystem for comprehensively probing cellular metabolism. Nature Communications, 10(1):1–13, 2019.

[9] M. Mori, T. Hwa, O.C. Martin, A. De Martino, and E. Marinari. Constrained allocation flux balance analysis. PLoS computational biology, 12(6):e1004913, 2016.

[10] Y. Nesterov. Introductory lectures on convex optimization: a basic course. Kluwer Academic Publishers, 2004.

[11] E.J. O’Brien, J.A. Lerman, R.L. Chang, D.R. Hyduke, and B.Ø. Palsson. Genome-scale models of metabolism and gene expression extend and refine growth phenotype prediction. Molecular systems biology, 9(1), 2013.

[12] A.-M. Reimers, H. Knoop, A. Bockmayr, and R. Steuer. Cellular trade-offs and optimal resource allocation during cyanobacterial diurnal growth. Proceedings of the National Academy of Sciences, 114(31):E6457–E6465, 2017.

[13] A. Schmidt, K. Kochanowski, S. Vedelaar, E. Ahrné, B. Volkmer, L. Callipo, K. Knoops, M. Bauer, R. Aebersold, and M. Heinemann. The quantitative and condition-dependent *Escherichia coli* proteome. Nature biotechnology, 34(1):104, 2016.

[14] L. Tournier, A. Goelzer, and V. Fromion. Optimal resource allocation enables mathematical exploration of microbial metabolic configurations. Journal of mathematical biology, 2017.

[15] B.R.B.H. Van Rijsewijk, A. Nanchen, S. Nallet, R.J. Kleijn, and U. Sauer. Large-scale 13c-flux analysis reveals distinct transcriptional control of respiratory and fermentative metabolism in *Escherichia coli*. Molecular systems biology, 7(1), 2011.

